# Scalable Fabrication of 3D Structured Microparticles Using Induced Phase Separation

**DOI:** 10.1101/2021.07.14.451688

**Authors:** Sohyung Lee, Joseph de Rutte, Robert Dimatteo, Doyeon Koo, Dino Di Carlo

## Abstract

Microparticles with defined shapes and spatial chemical modification can enable new opportunities to interface with cells and tissues at the cellular scale. However, conventional methods to fabricate shaped microparticles have trade-offs between the throughput of manufacture and precision of particle shape and chemical functionalization. Here, we achieved scalable production of hydrogel microparticles at rates of greater than 40 million/hour with localized surface chemistry using a parallelized step emulsification device and temperature-induced phase-separation. The approach harnesses a polymerizable polyethylene glycol (PEG) and gelatin aqueous-two phase system (ATPS) which conditionally phase separates within microfluidically-generated droplets. Following droplet formation, phase separation is induced and phase separated droplets are subsequently crosslinked to form uniform crescent and hollow shell particles with gelatin functionalization on the boundary of the cavity. The gelatin localization enabled deterministic cell loading in sub nanoliter-size crescent-shaped particles, which we refer to as nanovials, with cavity dimensions tuned to the size of cells. Loading on nanovials also imparted improved cell viability during analysis and sorting using standard fluorescence activated cell sorters, presumably by protecting cells from shear stress. This localization effect was further exploited to selectively functionalize capture antibodies to nanovial cavities enabling single-cell secretion assays with reduced cross-talk in a simplified format.

---

Microparticles with defined shapes and chemical modification promise to transform the way we interface with cells and tissues, acting as *in vitro* cell carriers that can tune biochemical and physical signals,^1^ scaffolds to promote the growth and infiltration of cells *in vivo* to regenerate tissue,^2, 3^ compartments for high-throughput single cell analysis,^4, 5^ and solid phases for barcoded molecular assays.^6, 7^ Numerous approaches to manufacture shaped hydrogel particles have been developed including wafer scale photolithography,^8^ microfluidic emulsion polymerization, and continuous and stop-flow lithographic methods.^9, 10^ These conventional methods have had trade-offs between the throughput of manufacture and precision of particle shape and chemical functionalization (Table S1).

Droplet microfluidics, where single, double, or aqueous two-phase emulsions have been employed to generate shaped microparticles,^11, 12^ has emerged as a powerful platform to produce uniform microparticles with different functions and properties. In particular, crescent-shaped microparticles^5^ or hollow shell particles^13^ produced by polymerizing precursors following aqueous two-phase separation (ATPS), possess a sub-nanoliter size cavity which can hold cells, and can template water in oil emulsions for performing single-cell and digital molecular assays. The current approach to manufacture these shaped particles, such as “nanovials”, requires precise injection of multiple polymer precursors into flow focusing microfluidic geometries, limiting scalability. Further, these previous approaches lacked robust approaches to spatially pattern different chemical functionalities, limiting the flexibility of the platform. For example, by locally patterning cell adhesive proteins to the cavity region of the nanovials, cells can be preferentially bound within the nanovials, protecting them from shear stress during more vigorous handling steps such as emulsification, fluorescent activated cell sorting (FACS), or delivery *in vivo* into tissue for therapeutic applications. Additionally, selective adhesion can direct loading to an adhesive region or cavity sized to the dimensions of a single cell,^14–16^ improving loading statistics beyond random Poisson processes. While more scalable manufacturing devices such as parallelized step emulsifiers and highly parallelized flow focusing devices have been used to dramatically enhance the production rate of spherical microparticles,^3, 17–19^ high-throughput production of shaped 3D-particles or capsules which require two phases has not been achieved.

In this study, we use a new induced-phase separation concept to overcome tradeoffs between particle complexity and fabrication throughput for the manufacture of microparticles with tunable localized surface chemistry and shape. We fabricate monodisperse 3D-axisymmetric particles with a chemically-functionalized cavity using a parallelized step emulsification device and temperature-induced phase separation. Harnessing the conditional phase separation of polyethylene glycol (PEG) and gelatin, photocrosslinkable PEG and gelatin ATPS droplets were generated and crosslinked with UV light to form uniform 3D axisymmetric particles with geometries dictated by the balance of interfacial tensions between the different phases (Figure 1).

**Figure 1.**
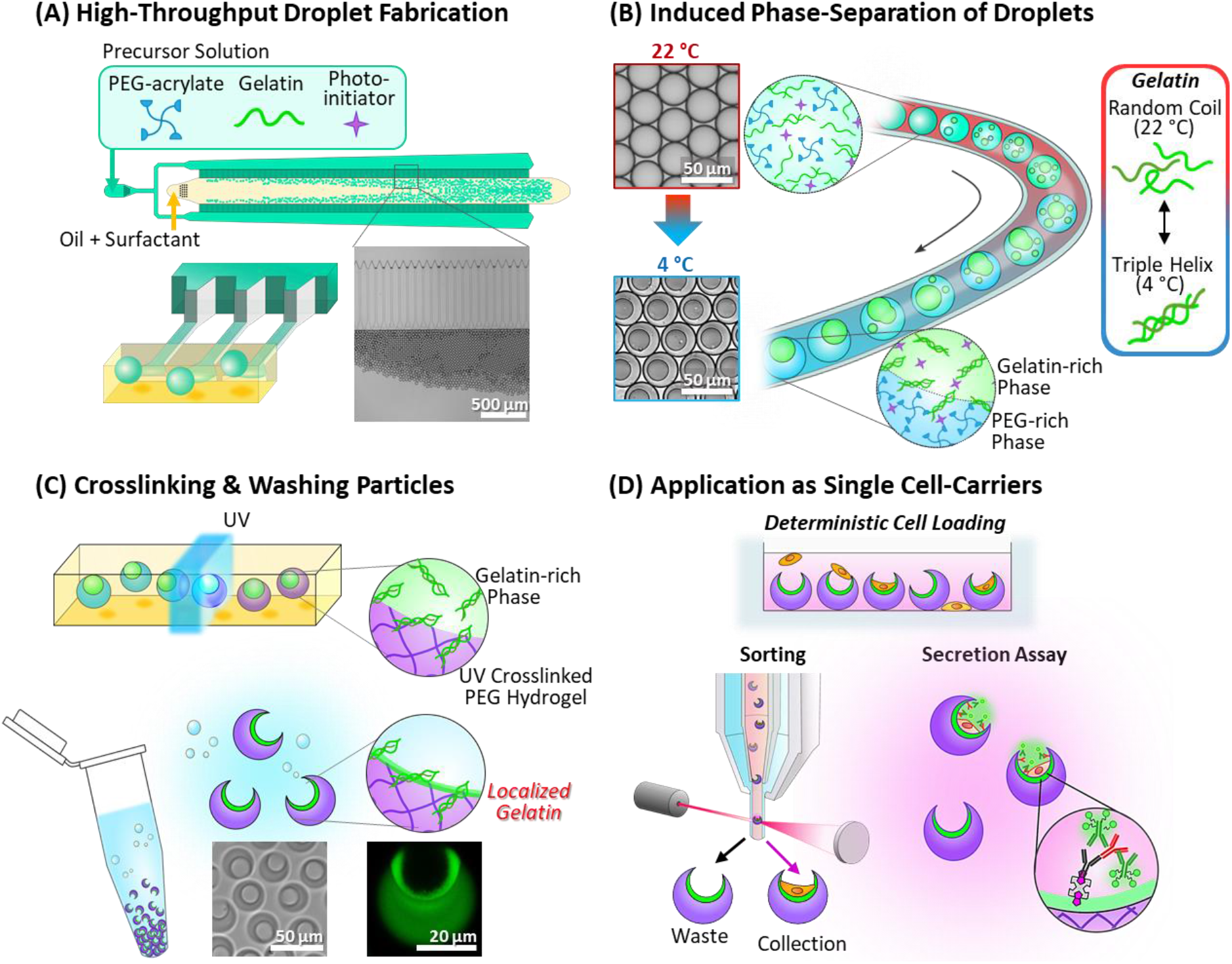
Overview of 3D structured microparticle fabrication using induced phase separation and their use. (A) Polymer precursors containing a mixture of PEG-acrylate, gelatin, and photo-initiator are injected into a high-throughput microfluidic droplet generator to create a uniform two-phase water in oil emulsion. (B) By reducing the temperature of the emulsions, PEG and gelatin undergo phase separation to create a three-phase PEG/gelatin/oil system. (C) The emulsion is exposed to UV light to selectively crosslink the PEG-rich phase and washed to recover 3D structure particles with gelatin remaining localized on the cavity surface. (D) The structured particles with localized surface chemistries act as cell carriers that protect cells from shear and show enhanced performance for single cell loading, secretion capture and live cell sorting using fluorescent activated cell sorters.

The engineered particles were found to be selectively functionalized on their inner cavities with higher densities of gelatin, which we showed was beneficial for their application as a platform for cell-carriers and reaction vessels to perform single-cell assays. The embedded gelatin promoted deterministic attachment of cells only within the cavities via integrin binding. The diameter and the opening of the crescent particles were also controlled to efficiently encapsulate single cells within nanovials, improving upon previous stochastic loading limitations governed by Poisson statistics. Cells adhered to nanovials could be sorted using standard FACS and viability was increased for cells attached to nanovials compared to unbound cells, suggesting these nanovials provide protection from fluid shear stresses during the sorting process. Finally, we showed the localized gelatin, when functionalized with capture antibodies, could be used to locally enrich secreted products from captured cells.

## RESULTS AND DISCUSSION

While PEG and dextran have been commonly used as two components of an ATPS in droplets,^5, 20, 21^ in this study, gelatin, instead of dextran, was used to enable us to trigger phase separation following massively parallel step emulsification using a temperature change. Gelatin, as denatured collagen, dissolves in water above its critical temperature and behaves as random coils in solution. Upon cooling gelatin molecules partially revert to triple helical collagen-like sequences altering the relative exposure of hydrophobic sites or other chemical groups on the surface of the gelatin molecules,^22^ which we found to affect the miscibility of gelatin with PEG. Thereby, we could control the miscibility and the partitioning behavior of PEG/gelatin solutions by varying the temperature and composition. Here, we exploit these properties to create multiphase water in oil templates that can be polymerized into 3D structured particles following UV polymerization.

Phase separation of PEG and gelatin is dependent on both the concentration of each component and the temperature of the system.^23^ Using a microfluidic droplet generator, we constructed two isothermal binodal curves, corresponding to which PEG and gelatin undergo phase transitions at 4 and 22 °C respectively (Figure 2A). We used gelatin derived from fish as it still remains liquid at 4 °C, allowing flow to form a minimal energy configuration unlike for porcine-derived gelatin. At concentrations above the binodal curves, the system undergoes phase separation to create PEG-rich and gelatin-rich regions within microscale water in oil droplets. For concentrations below the binodal curves, PEG and gelatin were miscible. The binodal boundary was found to be lowered by decreasing the temperature, which was attributed to favored interactions between gelatin molecules at lower temperatures (Figure 2B-C). By using compositions of the PEG/gelatin solutions located at points between the 4 and 22 °C binodal curves a transition is enabled from a miscible solution to a phase-separated state induced by the temperature change. This was confirmed for both bulk solutions (Figure S1) and in droplets (Figure 2C, S2).

**Figure 2.**
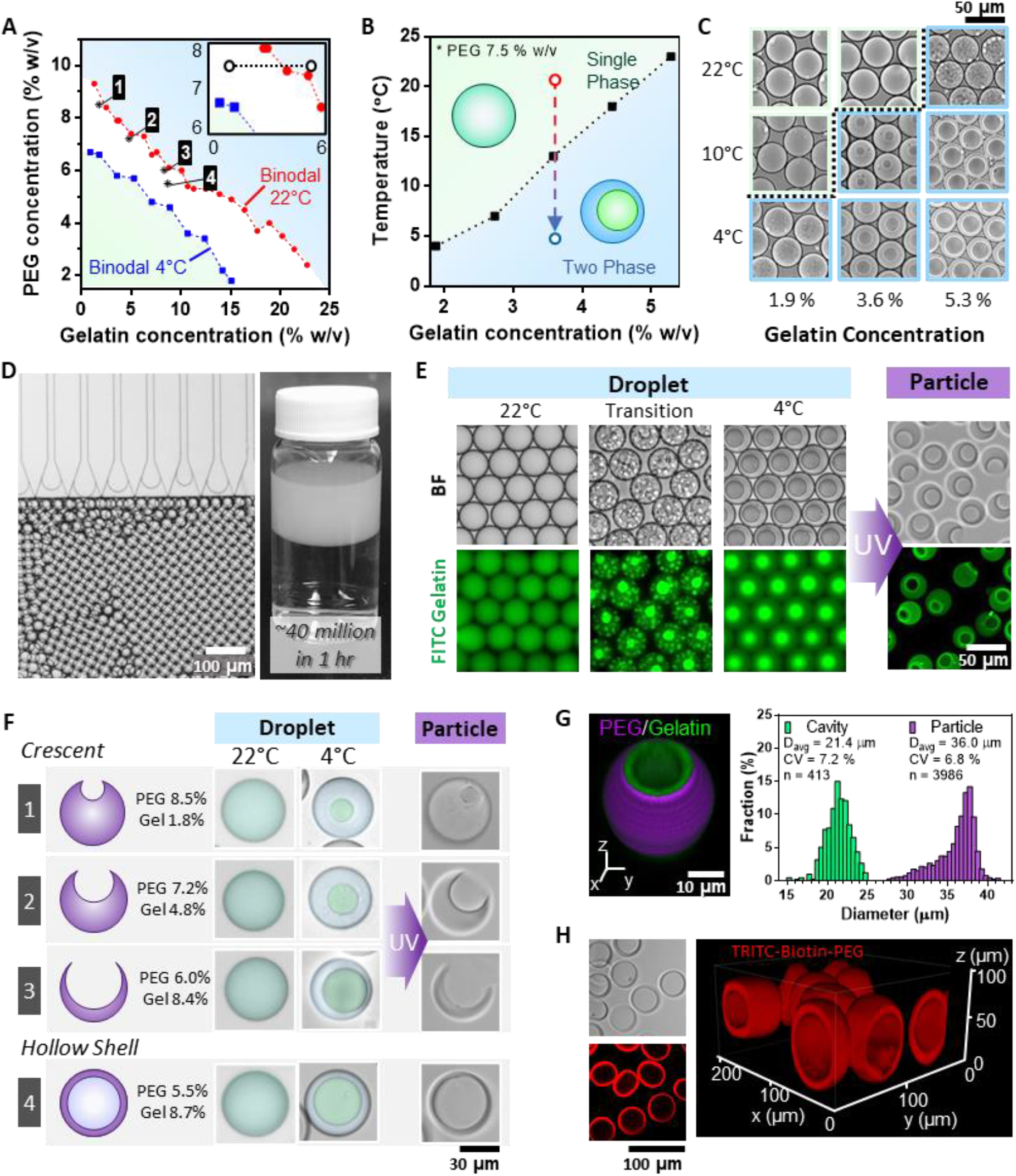
Fabrication of 3D structured particles using induced phase separation. (A) Phase diagram of PEG and gelatin. The isothermal binodal curve is shown for 22 °C and 4 °C. The dashed line in the inset shows the composition range whose transition temperature is indicated in Figure 2B. (B) At concentrations between the binodal curves, phase separation can be induced by adjusting the temperature of the system. (C) Example images of droplets at different gelatin compositions and temperatures (PEG concentration is 7.5% w/v). (D) Generation of unform single phase PEG/gelatin droplets using a highly-parallelized microfluidic droplet generator. (E) Microscopy images of PEG/gelatin (6.3 %w/v PEG and 4.5 % w/v gelatin) droplets undergoing induced phase separation from a reduction in temperature and resulting in monodisperse shaped particles after UV polymerization. Green fluorescent images show the distribution of FITC conjugated gelatin during the process to aid in visualization. (F) The structure of the resulting particles can be modified by adjusting the composition of PEG and gelatin (The corresponding compositions are shown on the plot in Figure 2A). Conditions are shown for crescent particles with different cavity ratios as well as fully enclosed hollow shell particles. Droplets are false colored to aid in visualization of PEG (green) and gelatin (blue) phases. (G, H) The morphology of the crescent shaped nanovials (G) and hollow shell particles (H) was confirmed using confocal microscopy.

We use a parallelized microfluidic droplet generator to fabricate structured microparticles in high-throughput by exploiting PEG/gelatin compositions that undergo induced phase separation (Figure S3A). Step emulsification devices are beneficial in that they can be easily scaled to generate droplets in high-throughput using 100s to 1000’s of parallel channels.^3, 17, 24^ However, they are limited in that only a single phase or stable mixture can be easily introduced when parallelized. Leveraging the PEG/gelatin ATPS system we are able to surpass these limitations by first generating droplets with a step emulsification device starting at a single phase composition between the binodal lines (Figure 2D, S3B Video S1), and then inducing phase separation by reducing temperature to create uniform multiphase geometries. As the temperature of the system is reduced very small domains of the gelatin-rich phase form within the larger drops, which coalesce and coarsen over time to form a single large spherical gelatin-rich domain in each drop (Figure 2E, S4 Video S2-4). The phase-separated droplets were then exposed to UV light to crosslink the PEG portion, yielding particles with crescent-shaped cross-sections having an average diameter of 36.0 μm (CV = 6.8 %) and average cavity diameter of 21.4 μm (CV = 7.2 %) (Figure 2G) after washing steps. Nanovial particles were generated at rates of 40 million/hour which is ~11 times faster than previous methods that used a single flow-focusing device (Figure S3C).^5^ Nanovials of various sizes could be generated by using step-emulsifiers with different channel heights (Figure S3D). The morphology of the ATPS droplets and resulting particles can be adjusted by changing the composition of PEG and gelatin. The compositions affect both the relative volumes of the PEG-rich and gelatin-rich regions as well as the balance of interfacial tensions between the PEG-rich, gelatin-rich, and oil phases. Compositions were found that resulted in repeatable fabrication of crescent particles with exposed cavities (nanovials). Increasing the concentration ratio of gelatin to PEG resulted in droplets with a higher volume fraction of the gelatin-rich phase, and nanovials with a larger exposed cavity when crosslinked (Figure 2F). Compositions were also found that result in particles with completely enclosed cavities (i.e. hollow shell particles) (Figure 2H).

Our unique fabrication approach results in a localization of gelatin on the inner cavity surface of the crescent shaped particles, which is advantageous for cell microcarrier applications utilizing this nanovial geometry. We found that during our particle manufacturing process gelatin molecules near the interface of the PEG-rich and gelatin-rich phases become trapped in the cross-linked surface. We used fluorescein isothiocyanate (FITC)-conjugated gelatin to visualize this localization effect using fluorescence and confocal microscopy (Figure 2E-G, 3A, S5, Video S5). We found that even after vigorous washing steps that the gelatin remained embedded in the particle surface indicating either physical entanglement and/or chemical bonding had occurred.

**Figure 3.**
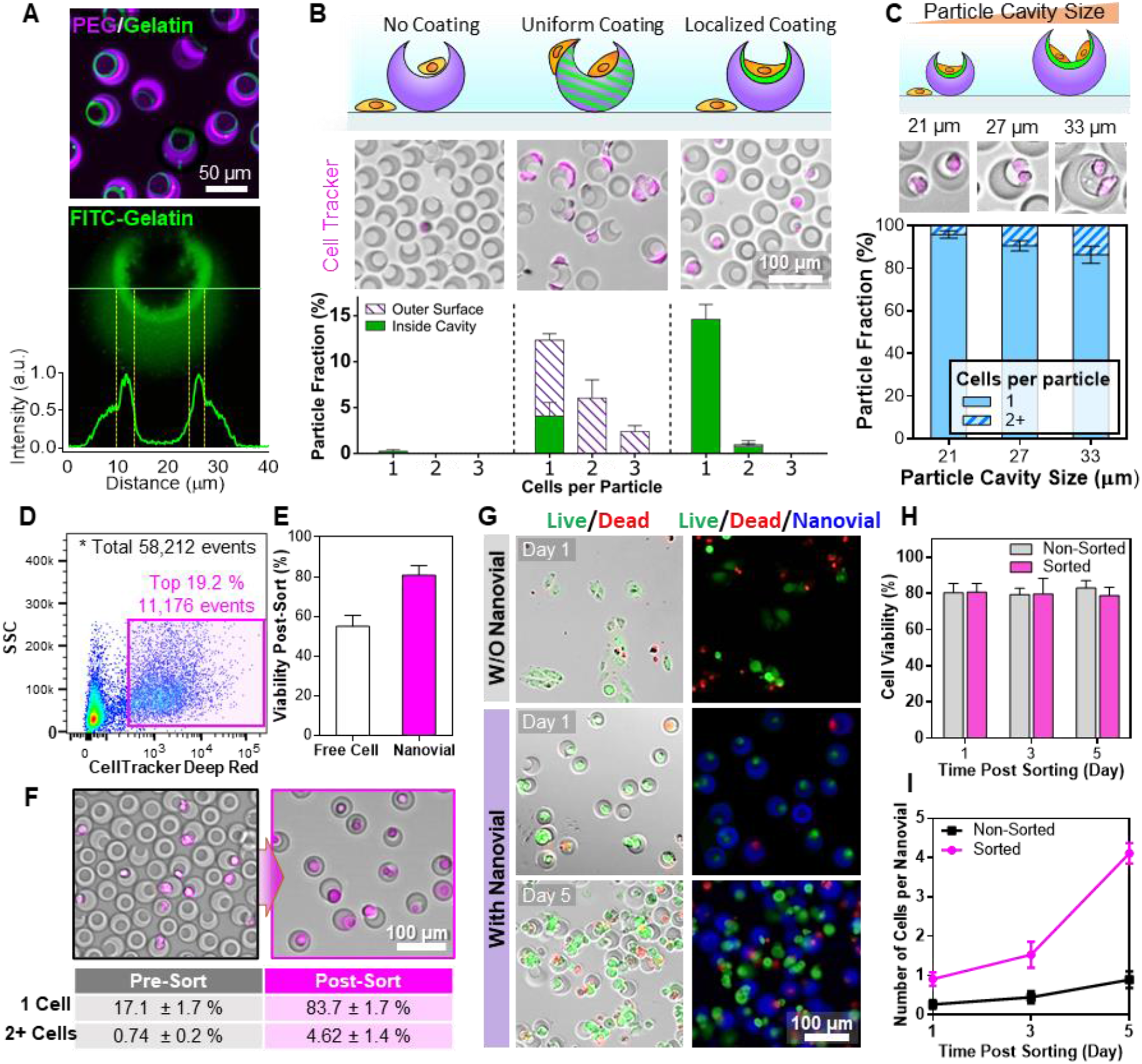
Nanovials with gelatin coated cavities for cell loading and sorting. (A) Confocal microscopy shows the localization of gelatin on the inner cavity surface of the crescent shaped particles. (B) Cell loading efficiency on nanovials with different distributions of binding moieties (No coating – PEG particles without RGD or gelatin, Uniform coating – PEG particles with RGD uniformly distributed, Localized coating – gelatin localized to the inner cavity). (C) The fraction of nanovials with single cells increases as the particle cavity size approaches the cell diameter. (D) Sorting nanovials loaded with cells based on CellTracker signal using FACS. (E)Viability of suspended cells (Cell) and cells loaded in gelatin-nanovial cavities (Nanovial) after sorting. Cells bound to nanovials showed significantly higher cell viability following sorting, suggesting these nanovials provide protection from fluid shear stresses during the sorting process (****p < 0.0001). (F) Cell-laden nanovials were sorted with high efficiency using FACS. (G) Example images of live/dead stained cells after sorting that were either freely suspended or bound to nanovials during the sorting. (H) Viability of cells loaded on nanovials remained ~80% over 5 days of culture for both non-sorted and sorted samples. (I) Average number of cells in nanovials increases as they proliferated for both sorted and non-sorted samples.

To investigate the effect of the localized gelatin layer on cell adhesion and growth in nanovials, we compared the cell loading on three different particles with no binding motif, with uniform RGD motifs, and with localized gelatin, respectively (Figure 3B). When Chinese hamster ovary (CHO) cells were seeded at a 0.6:1 cell-to-particle ratio, the particles without cell binding motifs had few cells attached to the particles (0.26 ± 0.17 % of particles had adhered cells). More cells were bound to RGD-coated nanovials as expected (20.73 ± 1.77 % of particles), but a significant fraction of cells was bound to the outside of the particle (80.31 % of cells bound to particles). Particles with localized gelatin showed a significantly larger fraction of cells bound to the particle cavity, 15.39 ± 1.62 % of particles, while less than 1.33 ± 0.73 % of them were bound to the outer surface (a 60-fold improvement as compared to uniformly coated particles). By avoiding external adhesion, the majority of cells can be localized to cavity region which can help ensure loading of single cells as well as reduce unwanted shear stress during handling steps, improving cell viability.

Localized adhesion combined with size exclusion effects of the cavity enables deterministic loading of single cells into the particle cavities. We found that for the same cell seeding concentrations, nanovials with uniformly distributed binding moieties yielded a significantly larger fraction of nanovials containing more than one cell (> 80 % cell containing nanovials) than nanovials with gelatin localized to the cavity (~ 1% of cell containing nanovials). The high-multiplet fraction for the nanovials with uniformly distributed binding moieties was attributed to the larger fraction of cells bound to the outside of the particles and resulted in loading statistics slightly worse than Poisson loading (Figure S6A). For nanovials with localized gelatin, the reduction in cells binding to the outer surface combined with exclusion effects of the inner cavity size was found to improve loading of single cells beyond distributions predicted by Poisson statistics. Both cells loaded in RGD-coated nanovials and localized gelatin-nanovials showed high viability (>80 % over 5 days of culture (Figure S7)). Testing a range of nanovial sizes, we found that as the cavity proached the average size of the cells (~17 μm diameter) the fraction of nanovials with singlets increased and multiplets decreased (Figure 3C, S6B). This effect became more evident at the higher cell seeding densities where the fraction of multiplets was reduced by as much as 61% compared with Poisson loading (Figure S6C). Although there is a tradeoff with loading efficiency for smaller cavity sizes, as it is more difficult for cells to fall into the cavity opening, we expect that improved singlet loading can dramatically improve utility of the cell carriers for applications where clonality is critical and overcomes one of the long-standing limitations of microfluidic platforms that are dictated by Poisson loading.^25^

The 3D-structured nanovials with gelatin functionalized cavities facilitate cell growth and prevent cell death during standard assays that can induce high fluid dynamic shear stress such as fluorescence activated cell sorting (FACS). We sorted both freely suspended cells and cells adhered in nanovial cavities at high-throughput using FACS (~270 events/second) (Figure 3F) and expanded cells after sorting over several days (Figure 3G). We found that cells bound in the nanovial cavities showed significantly higher viability than unbound cells after sorting (54.9% vs. 80.0%, p < 0.0001) (Figure 3E, S8). We found that there was no significant difference in viability of nanovial bound cells that were sorted vs samples that were not sorted (Figure S8). After sorting, the cells loaded in nanovials proliferated well over 5 days of culture and there were no significant differences in viability as compared to the non-sorted control group (Figure 3H-I). We attribute the improved viability to the cavity acting as a protective shelter that reduces hydrodynamic shear stress on cells during the sorting process.^26^ This improved viability is critical for improving the success of clonal expansion after sorting single cells and could be advantageous for analysis and recovery of adherent cells.

Gelatin localized on the inner particle surface enables facile spatial modification of particles with other biomolecules of interest. Due to the abundance of functional handles such as free amines and carboxylic acid, gelatin is a convenient base for bioconjugation. For example, free amines can be easily linked to using N-hydroxysuccinimide (NHS) ester conjugates (Figure 4A). Using an NHS-biotin conjugate we selectively modified the inner cavity of the particles with biotin, a biomolecule commonly used as a high affinity linker for antibodies, proteins, or oligonucleotides via the extremely high affinity biotin-streptavidin non-covalent interaction. To visualize the difference in localization we fabricated particles with both biotin incorporated throughout the PEG backbone (Biotin-PEG) and biotin linked directly to the localized gelatin (Biotin-Gelatin) and stained with fluorescent streptavidin. Fluorescence and confocal microscopy revealed that the Biotin-PEG particles have a uniform distribution of biotin groups, while Biotin-Gelatin particles have a significant increase in fluorescence intensity around the inner surface of the cavity indicating a higher concentration of available biotin groups (Figure 4B-C).

**Figure 4.**
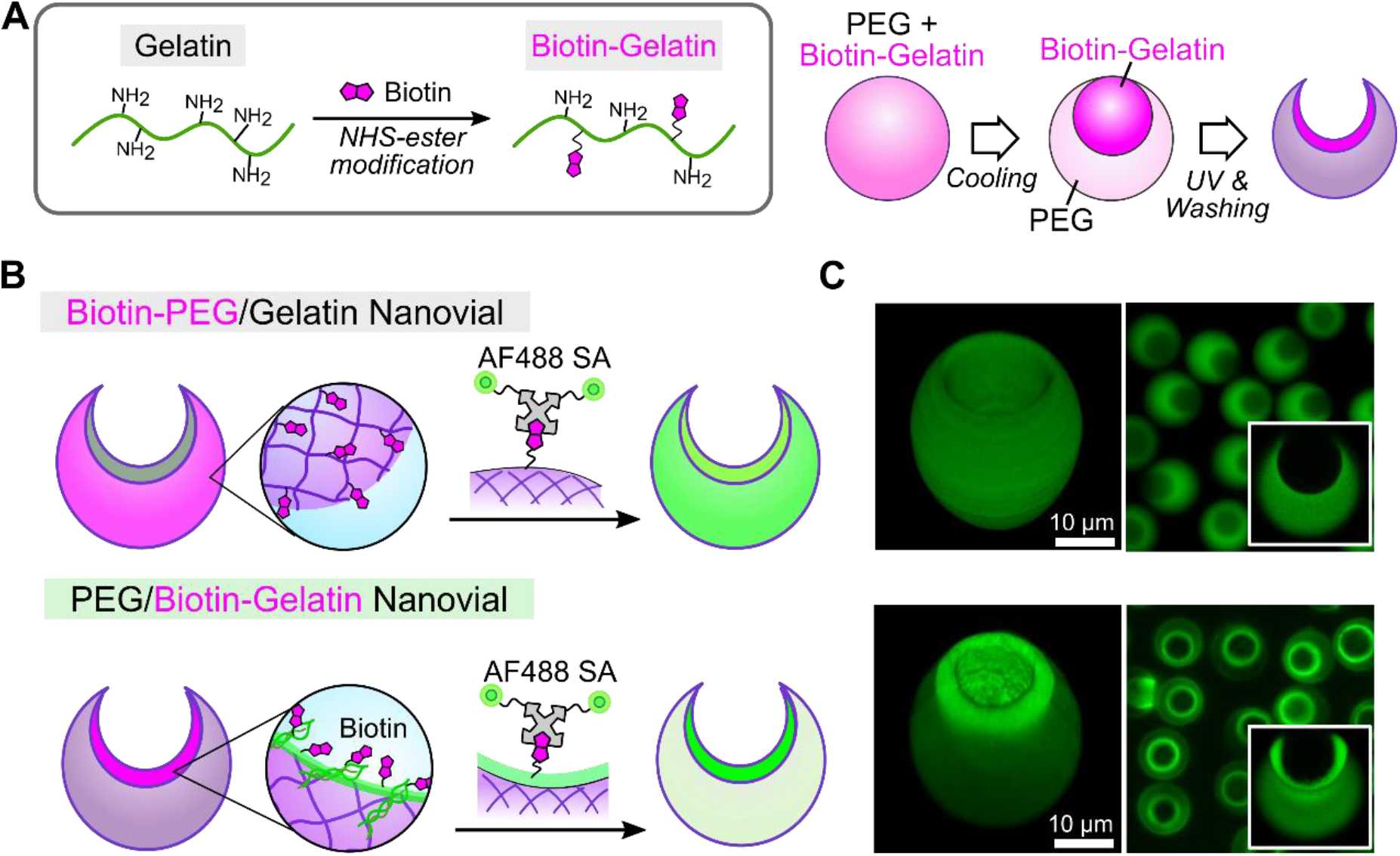
Spatial modification of particles with biomolecules using localized gelatin. (A) Free amine groups on gelatin are conjugated with biotin using NHS-ester modification. (B) Both biotin modified PEG (Biotin-PEG) and biotin modified gelatin (Biotin-Gelatin) are conjugated with AlexaFluor™ 488 conjugated streptavidin after fabrication. (C) Fluorescence and confocal imaging show increased fluorescence intensity in the inner cavity of Biotin-Gelatin nanovials indicating localization of biotin.

The ability to spatially pattern biomolecules can be exploited to enhance detection accuracy for high throughput analysis of single cell secreted products. Previously, we have shown that nanovials can be used to perform single cell secretion assays by using the particle surface to immobilize secreted antibodies and perform a fluorescent sandwich immunoassay. To reduce cross-talk between cells and ensure accurate measurements an emulsification step was required with the previous particles.^5^ Here we show that performing the assay with nanovials in which biotin is localized to the inner cavity surface substantially reduces the amount of cross-talk between cells enabling single cell secretion assays, without the need for extra steps to prevent cross-talk.

To characterize the effect of localized biotin on cross-talk, we measured the secretion of a Human immunoglobin G (IgG) against interleukin 8 (IL-8) produced by CHO cells loaded on both Biotin-PEG nanovials and Biotin-Gelatin nanovials (Figure 5A). We characterized the amount of cross-talk by introducing a population of empty control particles with a unique fluorescent label prior to the incubation step and measured the relative amount of secretions that were bound to these control “empty nanovials” compared to the cell containing population by flow cytometry (Figure 5B-C, S9).

**Figure 5.**
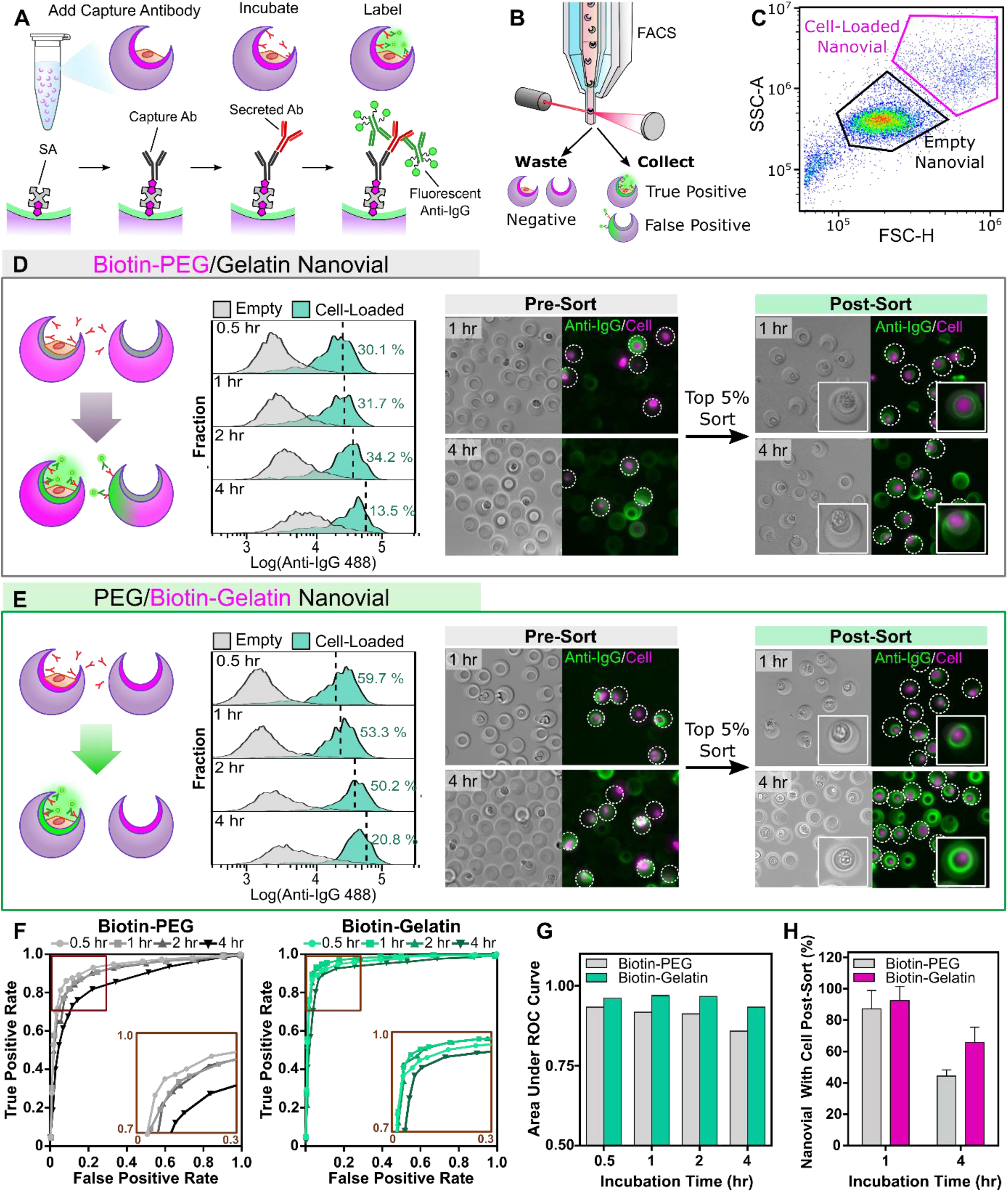
Localized conjugation of affinity agents reduces cross-talk for single cell secretion assays. (A) Single cell secretion assays were performed by first loading human IgG producing CHO cells into nanovials. After cell adhesion free biotin groups were conjugated with streptavidin and biotinylated antibodies against IgG. Cells were then incubated for different durations to accumulate secreted molecules on the surface of the particles and captured secretions were then fluorescently labeled with fluorophore conjugated antibodies. (B) The nanovials were analyzed and sorted in high-throughput based on secretion signals using FACS. (C) A flow cytometry scatter plot of 0.5 hr incubated nanovials, showing distinct scatter signal for nanovials containing cells. (D,E) Flow cytometry analysis and microscopy images showed that assays with Biotin-Gelatin nanovials (E) led to higher secretion signal and lower background intensity on empty nanovials as compared to Biotin-PEG nanovials (D) due to the localized capture antibody in cavity. The dashed lines in the histograms show the threshold to exclude the bottom 99% of control empty nanovials. The samples were stained with CellTracker before imaging to aid in visualization of cells. (F) ROC analysis was performed to compare the classification accuracy for Biotin-PEG and Biotin-Gelatin nanovial-based assays. (G) Area under the ROC curve indicates that Biotin-Gelatin nanovials enable more accurate classification than Biotin-PEG nanovials due to reduced cross-talk (p < 0.1) across all incubation times. (H) Nanovial fraction containing cells after incubating for 1 and 4 hours and sorting, reflecting the cross-talk differences between Biotin-PEG and Biotin-Gelatin.

We found that Biotin-Gelatin nanovials possessed a higher secretion signal and lower background intensity as compared to Biotin-PEG nanovials, indicating that the localized capture antibody in cavity of nanovials enriched the secretion signals and reduced secretion leak from cells to neighboring empty nanovials (Figure S10A). We defined a threshold of fluorescence intensity to exclude the bottom 99% of control nanovials (1 % false positive rate) and identified the percent of cell-loaded nanovials (true positive rate) that were above this threshold (Figure 5D-E). For Biotin-PEG nanovials we found that ~30-34% of cell-loaded nanovials possessed positive signal above this threshold when incubated within 2 hrs, which decreased substantially by 4 hrs to ~13% due to an increase in crosstalk to control nanovials (Figure 5D, S10B). For the Biotin-Gelatin particles, ~60% of cell-loaded nanovials were detectable above threshold after 30 minutes of incubation, which decreased slowly over time to ~21% at 4 hrs, indicating a significant reduction in cross-talk as compared to Biotin-PEG nanovials (Figure 5E, S10B).

To more fully characterize the capability of selecting out cell-containing nanovials from nanovials with signal cross-talk we performed receiver operating characteristic (ROC) analysis, a standard method to assess classification accuracy independent of a single threshold. For each condition, a curve of true positive rate versus its false positive specificity were obtained across a series of cutoff levels to depict the trade-off between the sensitivity and specificity (Figure 5F) and the area under the curve (AUC) for each ROC curve was calculated (Figure 5G). The ROC curve indicated that Biotin-Gelatin nanovials overall provide higher accuracy than Biotin-PEG nanovials with a minimum trade-off between true positive rate and false positive rate (Figure 5F). While both nanovials allow accurate detection of secretions (AUC > 0.85), Biotin-Gelatin nanovials showed significantly higher AUC values (AUC > 0.93) than Biotin-PEG nanovials due to the reduced cross-talk (p < 0.1) (Figure 5G). This was also reflected in fluorescence microscopy images of the samples (Figure S11). The reduction in cross-talk for Biotin-Gelatin nanovials was further evidenced by inspecting nanovials sorted based on the top 5% of fluorescence intensity. For shorter incubation times (1 hr) both the Biotin-PEG and Biotin-Gelatin particles had a relatively low fractions of nanovials without attached cells: 7.1% and 12.1% respectively (Figure 5D-E, H). Note that these empty nanovials also likely include some amount of nanovials in which cells may be dislodged during processing and sorting. After 4 hours the amount of empty nanovials sorted for the Biotin-PEG nanovial condition increased to 55.5 ± 4.0 % while Biotin-Gelatin nanovial sorts yielded only 33.8 ± 8.6 % empty nanovials (Figure 5H). We hypothesize that having the antibody binding sites localized in the cavity reduces the amount of leaked secretions reaching the binding sites of other particles from convective transport. Convective flows into nanovial cavities are expected to be reduced given the boundary conditions, therefore requiring slower diffusive transport to reach the reactive surfaces in the cavities. Further, it is possible that the higher concentration of binding sites in the cavity of the Biotin-Gelatin particles more efficiently captures secretions before they have time to diffuse out of the cavity or be convected away, further reducing crosstalk. Taken together, we conclude that locally functionalized nanovials can effectively capture the secretions from the single cells with lower cross-talk, enabling more accurate analysis and sorting of desired sub-populations with a reduced number of processing steps than previous approaches.

## CONCLUSION

To summarize, this study presents a scalable approach to fabricate 3D structured microparticles that can be locally functionalized with biomolecules of interest. Our unique temperature sensitive PEG/gelatin ATPS system allows for generation of multiphase droplets using any droplet generation system. Here we demonstrate compatibility with scalable step-emulsification devices to produce 3D structured particles at rates of 40 million/hr. Increased production to >1 billion/hr can easily be achieved by further scaling these devices,^27^ and can be potentially scaled further to >1 trillion/hr using other microfluidic approaches.^17^ We further demonstrate that crescent-shaped nanovial particles fabricated with this approach have an added benefit of gelatin localization inside the nanovial cavity. By exploiting this property, we show improved loading of cells into the particle cavities and demonstrate loading of single cells at rates that improve upon expectations based on Poisson statistics. The particle cavities further act as a shelter for the cells that protects them from shear stress during handling and processing by FACS, improving viability and promoting cellular growth. We further exploit this localization effect to selectively functionalize biotin and capture antibodies to the cavities of the particles enabling single-cell secretion assays with reduced crosstalk. This can be applied beyond producer cells to other cell types such as B cells and T cells where secretion profiling is of importance for the development of antibody and cell-based therapeutics.^28^

Beyond these initial demonstrations we anticipate broad impact of this technology across other systems. Cavity-containing cell carriers can be used more generally for improved viability of other adherent cell types sorted using FACS.^29^ Particles can be further modified to tune stiffness or extracellular matrix coatings to probe cellular response or provide more relevant cell microenvironments.^3^ Due to the scalability of our approach, it unlocks potential utility in other areas such as tissue engineering which typically requires a significantly larger amount of materials. For example, the protective cavity of the particles can be exploited to reduce the harmful effects of shear on in vivo delivery of cells for cell therapy applications while also providing a matrix for improved cellular growth.^30^ Localization of the binding moieties can further be exploited for the self-assembly of multicell systems.^31^ Aside from the crescent shaped nanovial particles we focus on in this work, this approach can also be applied for scaled fabrication of hollow-shell particles which can be used for studying clonal populations of mammalian cells, microalgae, and bacteria in biologically relevant environments,^13^ or act as an immunoprotective layer for allogeneic or xenogeneic cell therapies.

## EXPERIMENTAL SECTION

### Microfluidic Droplet Generator Fabrication

Step emulsification devices were fabricated as previously described.^3^ Master molds were fabricated on silicon wafers using a two-layer photolithography process to define the nozzle heights and reservoir heights. Devices were molded from the masters with PDMS and bonded to glass slides. Devices were treated with 2% trichloro (1H,1H,2H,2H-perfluorooctyl) silane (Sigma) in Novec 7500 (3M) to make the channel surfaces fluorophilic. Flow focusing devices used for the phase diagram studies were fabricated using a similar process.^5^

### Identifying Phase Separation Compositions in Bulk

For PEG and gelatin, 5000 Da 4 arm PEG acrylate (Advanced BioChemicals) and cold water fish gelatin (Sigma) were used. Nine different PEG/gelatin solutions comprising PEG at 5, 6, and 7 % w/v and gelatin at 5, 7.5, and 10 % w/v were prepared at room temperature and transferred to a refrigerator maintained at 4 °C. Three sets of conditions were identified in which a single phase of precursor materials transitioned to separated phases upon a temperature reduction from 22 °C to 4 °C.

### Identifying Phase Transition Temperature for Different Gelatin Concentrations

30 % w/v PEG, 20 % w/v gelatin and Dulbecco’s phosphate-buffered saline (DPBS), respectively, were injected into a flow focusing device with 3 aqueous inlets at different flowrates to precisely control the final compositions of water-in-oil droplets. As an oil phase, 0.5 % v/v Pico-Surf (Sphere Fluidics) in Novec 7500 was used. Five groups of droplets with fixed PEG concentration of 7.5 % w/v and different gelatin concentrations, 1.9, 2.7, 3.6, 4.4, and 5.3 % w/v, were generated. The PEG/gelatin droplets were collected in a downstream reservoir and immersed in a water bath in which the temperature was decreased from 22 °C to 4 °C by 1 °C every 30 minutes to identify at which temperature the droplets undergo phase separation.

### Binodal Construction at Two Different Temperatures

To build the binodal curve for PEG/gelatin droplets at 22 °C, different concentrations of PEG/gelatin droplets were generated using a flow focusing device as described above. Starting from 25 % w/v PEG, the target concentration of PEG was decreased by 0.75 % w/v and at each PEG concentration the minimum gelatin concentration required to yield phase separation in the ATPS droplet were measured. To build the binodal curve for PEG/gelatin droplets at 4 °C the above procedure used for 22 °C was followed but the generated droplets were collected in a downstream reservoir immersed in a 4 °C water bath.

### Fabrication of Nanovials with Localized Gelatin

For the dispersed phase, a homogeneous precursor solution with 6.3 % w/v PEG, 4.5 % w/v gelatin and 1.5 % w/v lithium phenyl-2,4,6-trimethylbenzoylphosphinate (LAP) dissolved in DPBS was injected into a parallelized step-emulsifier at 8 μL/min. In some cases, biotinylated PEG, biotinylated gelatin and FITC-gelatin were added to the solution to make Biotin-PEG nanovials, Biotin-Gelatin nanovials and FITC-labelled nanovials, respectively. The continuous phase comprised 2 % Pico-Surf in Novec 7500 injected at 16 μL/min. Single-phase PEG/gelatin droplets were generated and streamed through a Tygon tube (0.03” I.D., 0.0625” O.D, Murdock) immersed in a 4 °C water bath for temperature induced phase-separation of PEG and gelatin. The length of the tubing was adjusted to ensure full phase separation (~60 cm for 10 min incubation). The stream of phase separated droplets was directed into a PDMS reservoir submerged in the 4 °C water bath, and exposed to UV light (200 mW/cm^2^) for 1-3 seconds near the outlet region of the reservoir for polymerization. Upon UV exposure, the photocrosslinkable PEG components formed polymer networks while gelatin components remained unpolymerized. The crosslinked particles were collected, and the oil and the gelatin-rich drops were removed in a series of washing steps as previously described (for more details: see Methods in the Supporting Information),^5^ yielding crescent-shaped particles with localized gelatin in the surface of cavities.

### Fabrication of Nanovials with Uniform RGD Motifs

Nanovials with uniform RGD cell binding motifs were fabricated by injecting PEG and dextran solutions comprising RGD peptides into a flow-focusing device with 2 aqueous inlets as described in our previous study with some modification (for more details: see Methods in the Supporting Information).^5^

### Fabrication of Nanovials with No Binding Motif

Nanovials with no cell binding motifs were fabricated in the same way as RGD nanovials were made, except RGD was excluded from the composition.

### Labeling of Gelatin with FITC

50 mg gelatin was dissolved in 5 ml of pH 9.2 sodium carbonate/bicarbonate buffer. 0.25 mg of FITC was dissolved in 250 μL DMSO and slowly added to the gelatin solution. The reaction mixture was stirred for 6 hours in a 4 °C refrigerator. The reaction was quenched by the addition of 15 mg of ammonium chloride (50 mM) and the mixture was stirred for 2 more hours in the same condition. The solution was dialyzed against milli-q water with 14,000 dalton dialysis tubing for 5 days to remove excess FITC. The FITC-gelatin solution was collected in a 50 mL Falcon tube and stored in a −80 °C freezer and lyophilized.

### Biotinylation of Gelatin

10 mg of gelatin was dissolved in 100 μL of 3× PBS. The gelatin solution was mixed with 1 mg of N-hydroxysulfosuccinimide conjugated biotin (Thermo Sicentific, 21362) and incubated for 3 hours in room temperature. The solution was dialyzed against milli-q water using a dialysis device for > 24 hours and alioquoted into microtubes. The solutions were frozen at −80 °C and lyophilized.

### Characterizing Spatial Localization of Affinity Molecules

The localized gelatin in the nanovial was characterized using FITC-gelatin. PEG/gelatin nanovials were fabricated using a step-emulsifier as above, while 20 % of gelatin in the precursor solution was composed of FITC-gelatin. The nanovials were imaged in a green emission channel on a confocal fluorescence microscope (SP8-STED, Leica) and the intensity profile was obtained along a 2D slice across a nanovial revealing the stronger green signal along the surface of the cavity as compared to the rest of the nanovial. The localized biotin in Biotin-Gelatin nanovials was analyzed in the similar way. Biotin-Gelatin nanovials were fabricated with PEG/gelatin solution where biotinylated gelatin was used to constitue 20 % of the gelatin. The nanovials were stained by incubating for 30 minutes in 0.01mg/mL Alexa Fluor (AF) 488 conjugated streptavidin (Fisher Scientific) DPBS solution using the strong biotin-streptavidin affinity. After washing the nanovials three times, the fluorescent profile was analyzed using confocal microscopy.

### Cell Culture

CHO DP12 cells (ATCC CRL-12445) were cultured according to manufacture’s specifications. Briefly, cells were cultured in DMEM (Invitrogen) supplemented with 10 % fetal bovine serum (FBS, Invitrogen), 1 % penicillin/streptomycin (P/S, Invitrogen), 0.002 mg/ml recombinant human insulin (Sigma), 0.1% Trace Elements A (Fisher Scientific), 0.1% Trace Elements B (Fisher Scientific), and 200 nM Methotrexate (MTX, Sigma).

Seeding Cells on Nanovials: CHO cells were loaded by adding a cell solution to nanovials as described earlier with some modification (for more details: see Methods in the Supporting Information).^5^

### Cell Loading Statistics

Cell loading efficiency was evaluated using custom image analysis algorithms in MATLAB. After cell loading, nanovials were transferred to a well plate and imaged with a fluorescence microscope. The number of total particles were first identified using the particle fluorescence channel by the MATLAB script and the number of cell-laden nanovials were counted manually with detailed information including the number and location of cells within nanovials (n > 1500).

### Cell Viability Characterization

The viability of cells encapsulated in nanovials was evaluated using a live/dead assay. A staining solution of 2 μM calcein AM solution and 4 μM ethidium homodimer (EthD-1) was prepared in DPBS. Nanovials with cells were first concentrated in a conical tube by centrifugation and supernatant was aspirated. The concentrated nanovials were mixed with the staining solution and incubated for 30 minutes in a CO_2_ incubator. The samples were washed with DPBS, centrifuged and transferred to a well-plate for imaging. Live and dead cells were observed by a fluorescence optical microscope where living cells were detected by calcein AM (green fluorescence), and dead cells by EthD-1 (red fluorescence). The number of viable cells was quantified using ImageJ (NIH) software. Then, the viability rate was obtained by comparing the number of viable cells with total number of cells.

### Cell Viability Before and After Sorting

Cell viability of cells encapsulated in nanovials were characterized before and after sorting compared to unbound cells. CHO cells prestained with CellTracker were loaded in nanovials. Some of the samples were kept as non-sorted samples. The cell-loaded nanovials were sorted directly to a 96-well plate using a FACS machine (Sony SH800) based on the intensity of CellTracker signal at 100-500 events/second. Unbound cells were sorted in the same condition as a control. Viability of cells in different samples were assessed at days 1, 3, and 5 by staining live and dead cells as described above.

### Single Cell Secretion Assay and Cross-Talk Characterization

Nanovials were used as a secretion assay platform to capture human IgG targeting IL-8 produced by a CHO cell line (ATCC® CRL-12445™). Biotin modified nanovials were fabricated and CHO cells were seeded on nanovials at a cell-to-particle ratio of 0.8 as described above and cultured for 12 hours prior to the secretion test. Empty nanovials were stained with Alexa Fluor 350 streptavidin to distinguish them from the nanovials loaded with cells. The cell-loaded nanovials were mixed with the blue-stained empty nanovials as a method to characterize cross-talk. This was done in order to ensure signal on empty particles that was measured did not arise from cells that may have detached from the particles during various steps of the assay. All later steps were performed on the mixed nanovials. Mixed nanovials were diluted in a streptavidin solution (0.1 μg/mL streptavidin (Thermo Fisher, 434302) in washing buffer) at a 1:10 ratio and incubated for 10 minutes. The nanovials were washed with washing buffer three times. A capture antibody solution was prepared by adding 0.02 μg of biotin anti-FC (Thermo Fisher, A18821) per 1 mL of washing buffer. Nanovials were functionalized with capture antibody, biotin anti-FC (Thermo Fisher, A18821), by mixing the nanovial concentrate and the antibody solution at a 1:10 ratio and incubating for 10 min. The nanovials were washed three times and incubated in CHO cell media in a CO2 incubator for different durations 0.5, 1, 2, and 4 hours to allow cells to secrete IgGs which can bind to capture antibody. “Staining buffer” was made by mixing 10 % FBS, 1 % P/S and 0.05 % Pluronic F-127 in DPBS. The samples were washed three times with staining buffer and incubated in 0.01 μg/mL AF 488 secondary anti-human IgG (Invitrogen, A18821) solution for 30 minutes. The nanovials were washed three times with staining buffer and analyzed using a FACS machine at 100-500 events/second using both 350 nm and 488 nm laser excitation. For 1 and 4 hour incubated samples, nanovials with the top 5 % secretion signal were sorted directly into a 96-well plate pre-filled with culture media and imaged with a microscope. The nanovials before and after sorting were imaged with a fluorescence microscope (Nikon, Eclipse Ti-S). The samples were stained with CellTracker before imaging to aid in visualization of cells. The FCS data files from flow cytometry were analyzed using FlowJo and secretion signals from control and cell-loaded nanovials were compared.

### Receiver Operating Characteristic (ROC) Analysis

The signals from cell-loaded nanovials were identified based on their scatter readouts while control empty nanovials were found using their blue signals (Figure S9A). A threshold for anti-IgG signal was set and signals higher than the threshold from empty nanovials and cell-loaded nanovials were considered false positive and true positive signals, respectively. The true positive rates versus false positive rates at fifteen different cut-off levels were plotted and the area under the ROC curve was calculated using the trapezoidal method.

## Supporting information

Video S1 Droplet generation

Video S2 phase transition_BF x32

Video S3 phase transition_FITC x32

Video S4 Droplet downstream

Video S5. PEG-gelatin particle

## ASSOCIATED CONTENT

### Supporting Information

The following files are available free of charge.

Additional experimental details, materials, methods, tables and figures (PDF)

Video S1. Droplet generation (MP4)

Video S2. Phase transition_BF (×32) (MP4)

Video S3. phase transition_FITC (×32) (MP4)

Video S4. Droplet downstream (MP4)

Video S5. PEG/gelatin particle (PEG : purple, gelatin : green) (MP4)

## AUTHOR INFORMATION

### Author Contribution

S.L., J.D., and D.D. developed the idea. S.L. performed all experiments and data analysis. J.D. made the step-emulsifying microfluidic device. R.D. contributed to FACS cell sorting experiments. D.K. contributed to cell loading experiments. S.L, J.D., and D.D discussed the data and wrote the manuscript with input from all authors. D.D. supervised the study.

### Notes

The authors declare no competing financial interest.

## ACKNOWLEDGMENT

Confocal laser scanning microscopy was performed at the Advanced Light Microscopy /Spectroscopy Laboratory and the Leica Microsystems Center of Excellence at the California NanoSystems Institute at UCLA with funding support from NIH Shared Instrumentation Grant S10OD025017 and NSF Major Research Instrumentation grant CHE-0722519. We acknowledge support from the UCLA W. M. Keck Foundation COVID 19 Research Award Program.

