## Supporting Information for "Scalable Fabrication of 3D Structured Microparticles Using Induced Phase Separation"

**EXPERIMENTAL SECTION**

*Fabrication of Nanovials with Uniform RGD Motifs*: A PEG phase solution was prepared with 28.9 % w/w 10 kDa 4-arm PEG-Norbornene (Creative PEGWorks), 2 % w/w LAP, and 0.5 mg/ml Biotin-PEG-thiol (5000 MW, Nanocs) in DPBS. A dextran phase comprised of 11 % w/w 40 kDa dextran (Sigma), 0.7 % w/w dithiothreitol (DTT, Sigma), and 5 mM RGD peptide (Ac-RGDSPGERCGNH2, Genscript) in milli-q water. The PEG and dextran solutions were injected into a flow-focusing device at 0.5 µL/min separately. An oil phase comprised of 0.5% w/w Pico-Surf in Novec 7500 was injected at a rate of 10 µL/min to partition the aqueous phases into monodisperse water-in-oil droplets. The droplets were exposed to UV light (500 mW/cm$^2$)

for 1-2 seconds near the outlet region of the reservoir for polymerization. The oil and dextran-rich drops were removed using a series of washing steps.

*Washing Particles*: Surfactant oil was first removed by pipetting. A solution of 20% v/v perfluorooctanol (PFO, Sigma) in Novec 7500 and milli-q water were added consecutively to break the emulsions and transfer particles to the aqueous phase. Samples were centrifuged at 2500 ×g for 1 minute and oil phase was removed. Particles were washed with Novec 7500 twice to remove remaining PFO and surfactant. After removing the oil layer with pipetting, the residual oil was washed three times with hexane (Sigma). Samples were then washed three times with 70 % ethanol and then twice with milli-q water to remove gelatin. Particles were sterilized by incubating in 70 % ethanol overnight before use.

*Seeding Cells on Nanovials:* Nanovials were stained with AF 350 streptavidin (Fisher Scientific) before use. Nanovials were washed with cell culture media and concentrated in a conical. The amount of concentrated nanovials was calculated to cover the whole well area with a nanovial monolayer, which were 3.2, 4, and 4.8 $\mu L/cm^2$ for 36, 45 and 55 μm nanovials, respectively. The calculated amount of nanovials were transferred and dispersed in a well plate. CHO cells pre-stained with CellTracker deep red (Invitrogen, C34565) were seeded into the well with the nanovial monolayer at a target cell-to-nanovial ratio. The solution in the well was pipetted multiple times to evenly disperse the cells and particles across the well area. The sample was incubated for 2 hours in a $CO_2$ incubator to allow cells to adhere to nanovials. In order to remove unattached cells, samples were strained with a 20 μm reversible cell strainer (CellTrics) and washed with "washing buffer" comprised of 0.5 % bovine serum albumin (BSA), 1 % P/S and 0.05 % Pluronic F-127 in DPBS. Next, the cell strainer was flipped and washed with the washing buffer to recover the nanovials in a conical tube. The nanovials were centrifuged to remove washing buffer, resuspended in media, and cultured in a $CO_2$ incubator.

### SUPPORTING TABLE

| Approach | Shape Control | Multi material Patterning | Particle Size Range (μm) | Throughput [Particles/hour] | Material Efficiency(a) | Ease of Use | Demonstrated Bio Applications | Ref. |
|---|---|---|---|---|---|---|---|---|
| Stop Flow Lithography | 2D Mask | Yes | ~100 | 3.6X$10^{4}$ | <10% | **Med**<br>• Requires automated pumps/UV source<br>• Mask Alignment required | DNA oligomer detection | 1 |
| Scanning two-photon continuous flow lithography | 3D | Yes | ~50 | 3.6X$10^{4}$ | <1% | **Low**<br>• Requires specialized two photon excitation and scanning system | NA | 2 |
| Maskless Lithography | 2D | No | 60-180 | 6.0x$10^{4}$ | ~50% | **Med**<br>• Requires high-speed spatial light modulator | Multiplexed cell assays | 3 |
| Transient Liquid Molding | 3D, Intersecting 2D projections | Yes | 200 - 600 | 3X$10^{2}$ | <1% | **Low**<br>• Device manufactured for each design<br>• Requires automated pumps/UV source<br>• Mask Alignment required | NA | 4 |
| | 3D, Intersecting 2D projections | Yes | ~400 | 3X$10^{4}$ | <1% | | Cell microcarriers, cell viability assay, image flow cytometry | 5 |
| | 3D, Intersecting 2D projections | Yes | 200 - 600 | 3X$10^{4}$ | <1% | | Single cell secretion assay, amplified immunoassays | 6 |
| Droplet Polymerization | 1D, Spherical | No | 25 - 100 | 2X$10^{6}$ | 95% | **High**<br>• Uses common flow focusing device<br>• No UV or mask | Tissue engineering | 7 |
| | 1D, Spherical | No | 50 - 100 | 1.4X$10^{7}$ | >99% | **Very High**<br>• Compatible with any droplet generator<br>• No UV or mask | Tissue engineering | 8 |

| | | | | | | | | |
|---|---|---|---|---|---|---|---|---|
| 3D Co-Flow | 3D, Intersecting 2D projections | Yes | 100 - 400 | $2X10^4$ | 1-5% | **Med**<br>• 3D printed devices that can be outsourced<br>• Requires automated pumps/UV<br>• Mask Alignment required | Amplified immuno assays | 9<br>10 |
| ATPS<br>(PEGDA/DEX) | 3D revolved crescent | No | 20 - 80 | $1X10^6$ | ~95% | **High**<br>• Uses common flow focusing device<br>• Compatible with common UV sources<br>• No mask required | NA | 11 |
| | 3D revolved crescent | No | 125 - 200 | $1X10^4$ | ~95% | | Cell growth, cell killing assay | 12 |
| | 3D revolved crescent | Yes | 136 - 200 | $1X10^4$ | ~95% | | Cell killing assay | 13 |
| ATPS<br>(4APEGNB/DEX) | 3D revolved crescent | No | 35 – 85 | $4X10^6$ | ~95% | | Single-cell secretion assay, Flow cytometry, Digital PCR | 14 15 |
| ATPS<br>(4APEG/DEX) | 3D hollow shell | No | ~80-120 | $2X10^6$ | ~95% | **High**<br>• Uses common flow focusing device<br>• No UV or mask | Single-cell growth and productivity assay, Flow cytometry | 16 |
| ATPS<br>(4APEGA/Gelatin) | 3D, revolved crescent, hollow shell particle | Yes | 35 - 65 | $2X10^7$<br>($2X10^8$ – $1X10^9$ if scaled)(b) | ~99% | **Very High**<br>• Compatible with any droplet generator<br>• Compatible with common UV sources<br>• No Mask required | Single-cell secretion assay, Flow cytometry | This Work |

(a) Material Efficiency: Fraction of precursor material converted into particles
(b) Estimate ~$2X10^8$ $hr^{-1}$ throughput for 5000 channel step-emulsification device and ~$1X10^9$ $hr^{-1}$ throughput for parallelized flow focusing device

Table S1. Throughput comparison of devices used to fabricate microgels with different shapes.

#### SUPPORTING FIGURES

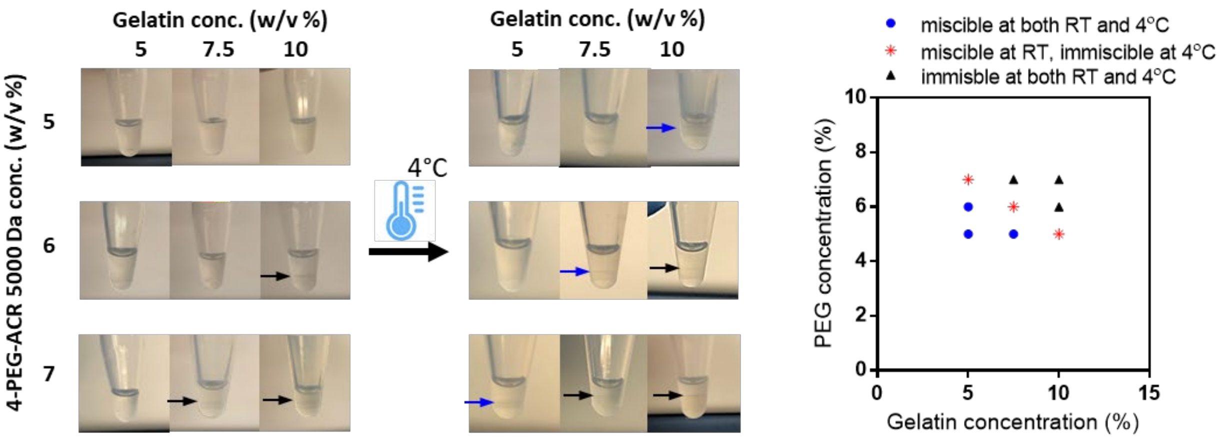


Figure S1. The phase separation behavior of a solution comprising different concentrations of 4 arm PEG acrylate 5000 Da and fish gelatin depending on temperature. Three sets of conditions are highlighted in red in which a single phase of precursor materials transitions to separated phases upon a temperature reduction from 25 °C to 4 ° C.

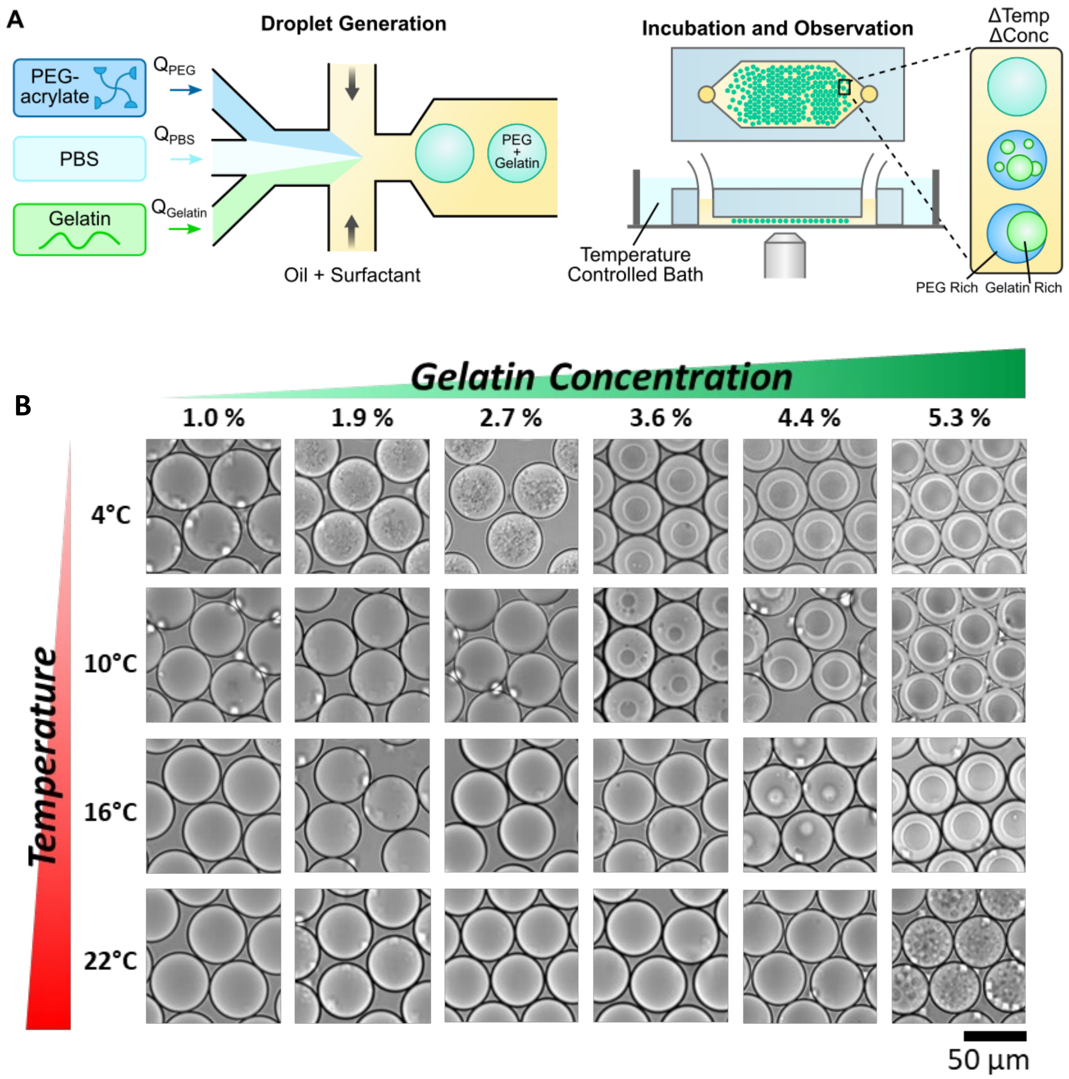


Figure S2. Microfluidic setup for construction of PEG-gelatin phase diagrams. (A) PEG acrylate, PBS, and gelatin are injected into separate inlets of a flow focusing device and mixed prior to a droplet generating junction. The composition of droplets is controlled by adjusting the corresponding flow rates. Droplets are collected into a PDMS reservoir in a temperature controlled bath and observed using an inverted microscope. (B) Example images of droplets with different gelatin compositions and temperatures (PEG concentration is 7.5% w/v). Phase separation is observed in several different conditions.

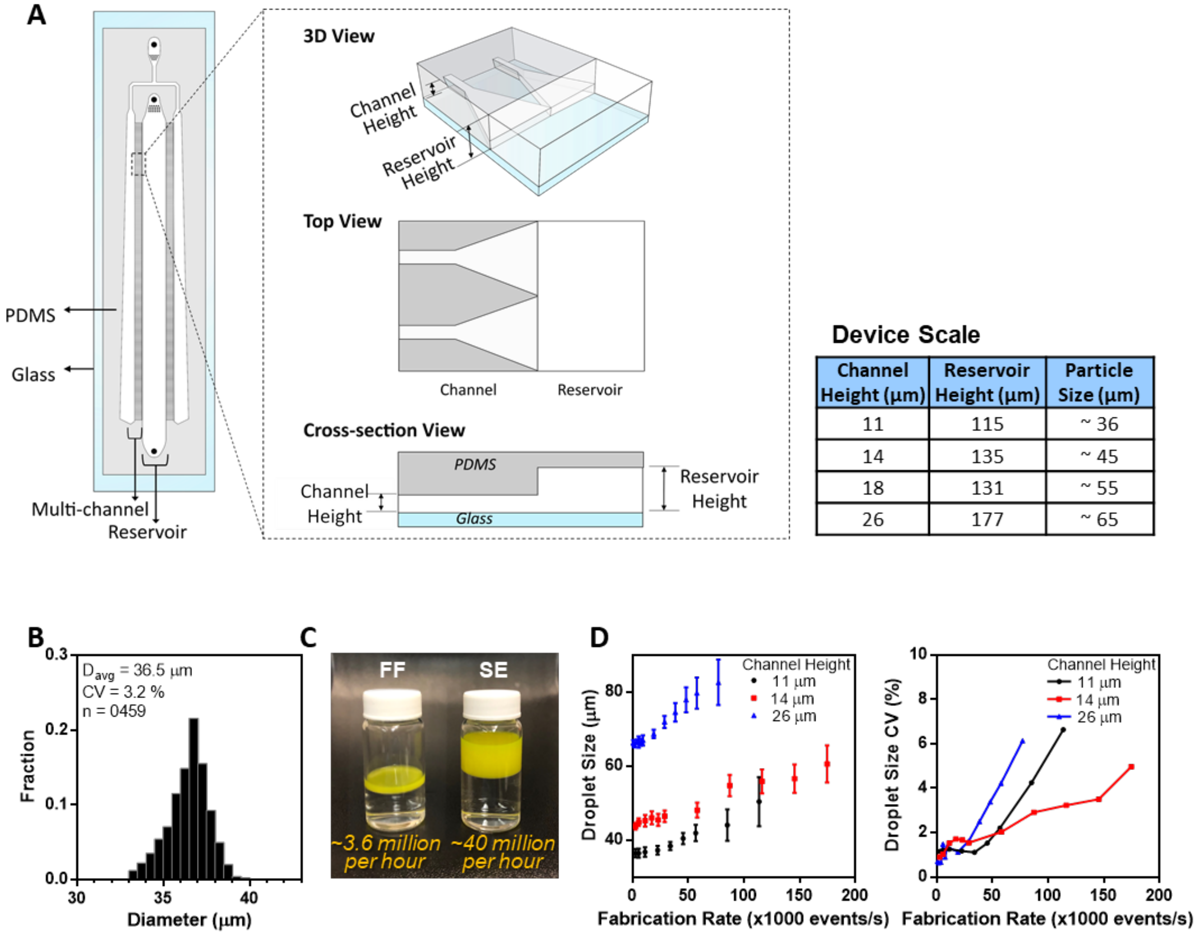


**Device Scale**

| Channel Height (μm) | Reservoir Height (μm) | Particle Size (μm) |
|---|---|---|
| 11 | 115 | ~ 36 |
| 14 | 135 | ~ 45 |
| 18 | 131 | ~ 55 |
| 26 | 177 | ~ 65 |

Figure S3. Characterization of uniform PEG/gelatin droplets generated using a highly-parallelized microfluidic droplet generator. (A) A schematic of a multi-channel step-emulsification microfluidic device which can be used for high-throughput production of droplets for induced phase separation-based manufacturing at high rates. (B) The step-emulsification device yielded droplets of average diameter 36.5 µm which were highly monodispersed (CV = 3.2 %) at a production rate of 1 mL/hr. (C) Visual comparison of the production rate of the parallelized step emulsification device versus a flow focusing device (1mL·h$^{-1}$ and 0.06 mL·h$^{-1}$, respectively). (D) Step-emulsifiers with channel heights of 11, 14, and 26 µm generated droplets with mean diameters of 36, 45, and 65 µm, respectively. Since the droplet formation in a step-emulsifier is mainly driven by interfacial tension,[17] the droplet size was determined by the height of the inlet channel and did not change significantly over a range of flow rates of both the dispersed miscible precursor solution and the continuous oil phases. However, as flow rate increases, collisions between the generated drops resulted in droplet coalescence, which can explain the increase in droplet sizes and their variance at higher flow rates, despite not reaching a jetting regime.

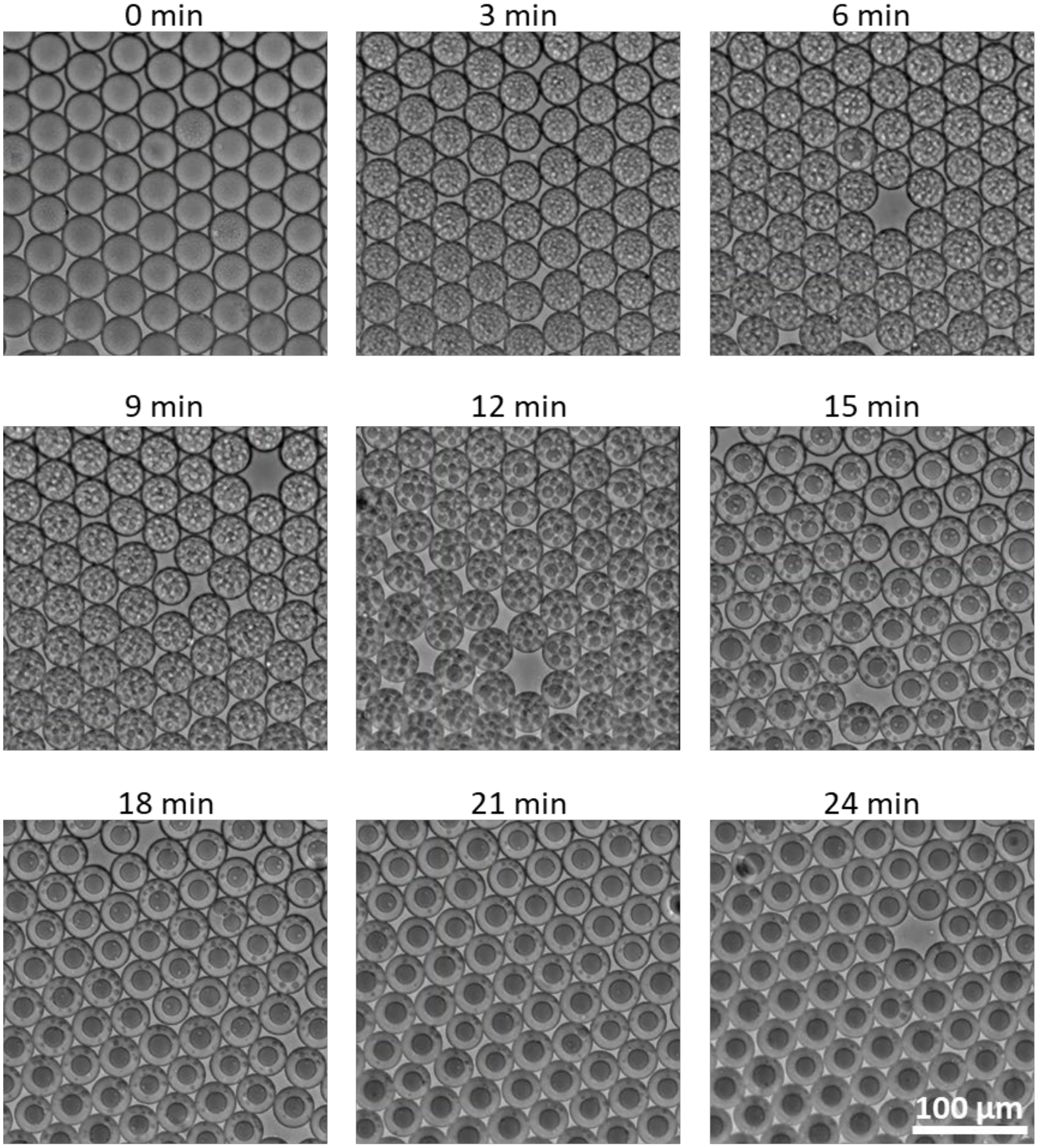


Figure S4. Timelapse images of PEG-Gelatin ATPS droplets undergoing induced phase separation. 45 µm droplets comprised of a precursor solution containing 6.3 % w/v PEG and 4.5 % w/v gelatin were first generated at room termparature. Droplets were imaged in a PDMS chamber and put on ice at the 0 min timepoint. Small partitions of gelatin were initially observed, and coarsened over time until a single PEG-rich and single gelatin-rich region remain at 24 min

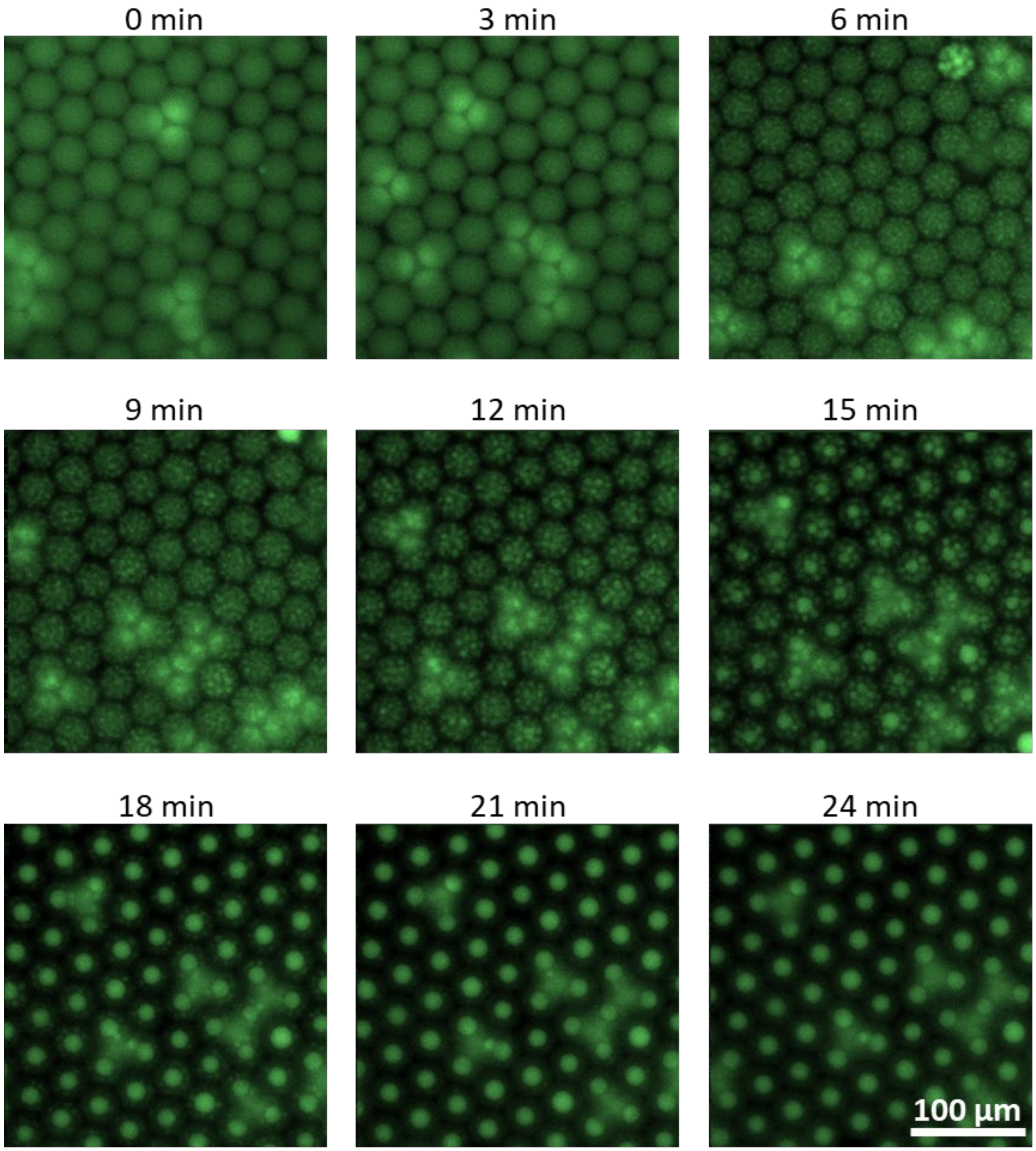


Figure S5. Fluorescence microscopy timelapse images of temperature induced phase separation for the same conditions described in Figure S5. Fluorescently tagged gelatin was added to the gelatin phase to visualize partioning behavior.

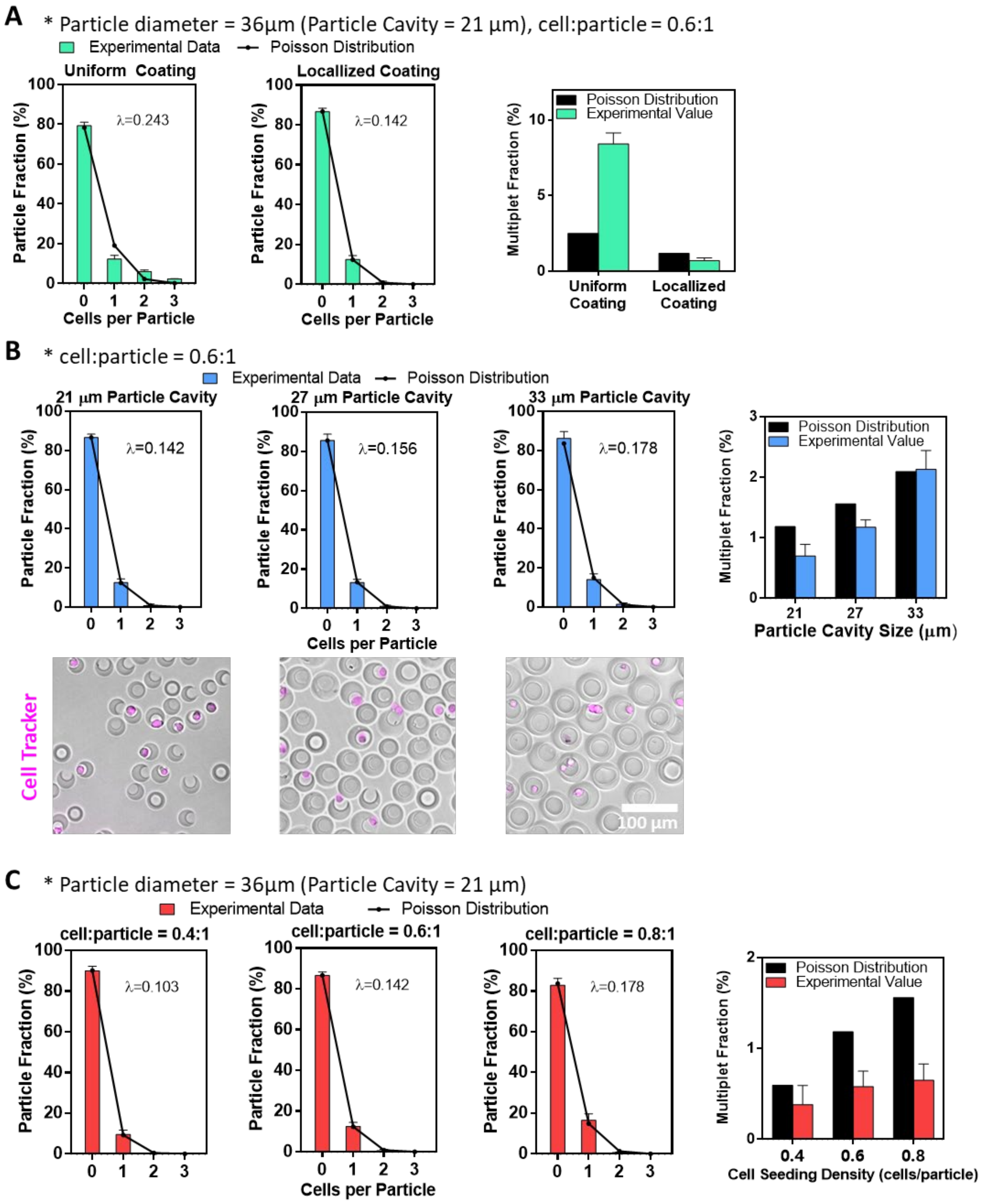


Figure S6. Further characterization of cell loading into nanovials in comparison to Poisson loading. (A) Comparison of loading into 36 micron nanovials with uniform binding moieties and localized binding moieties. Particles with uniform binding moieties have loading statistics worse than the Poisson distribution, likely due to cell clusters being able to bind to the outside of the nanovials. This trend is highlighted when looking at the fraction of nanvovials with multiplets (>1 cell per nanovial). Localized binding moieties in particles promotes deterministic cell loading and reduces the multiplet fraction below the number predicted by Poisson statistics. (B) As the cavity size approached the average size of the cells (~17 µm), the fraction of nanovials with singlets increased and multiplets decreased. (C) The deterministic loading became more evident at higher cell seeding densities.

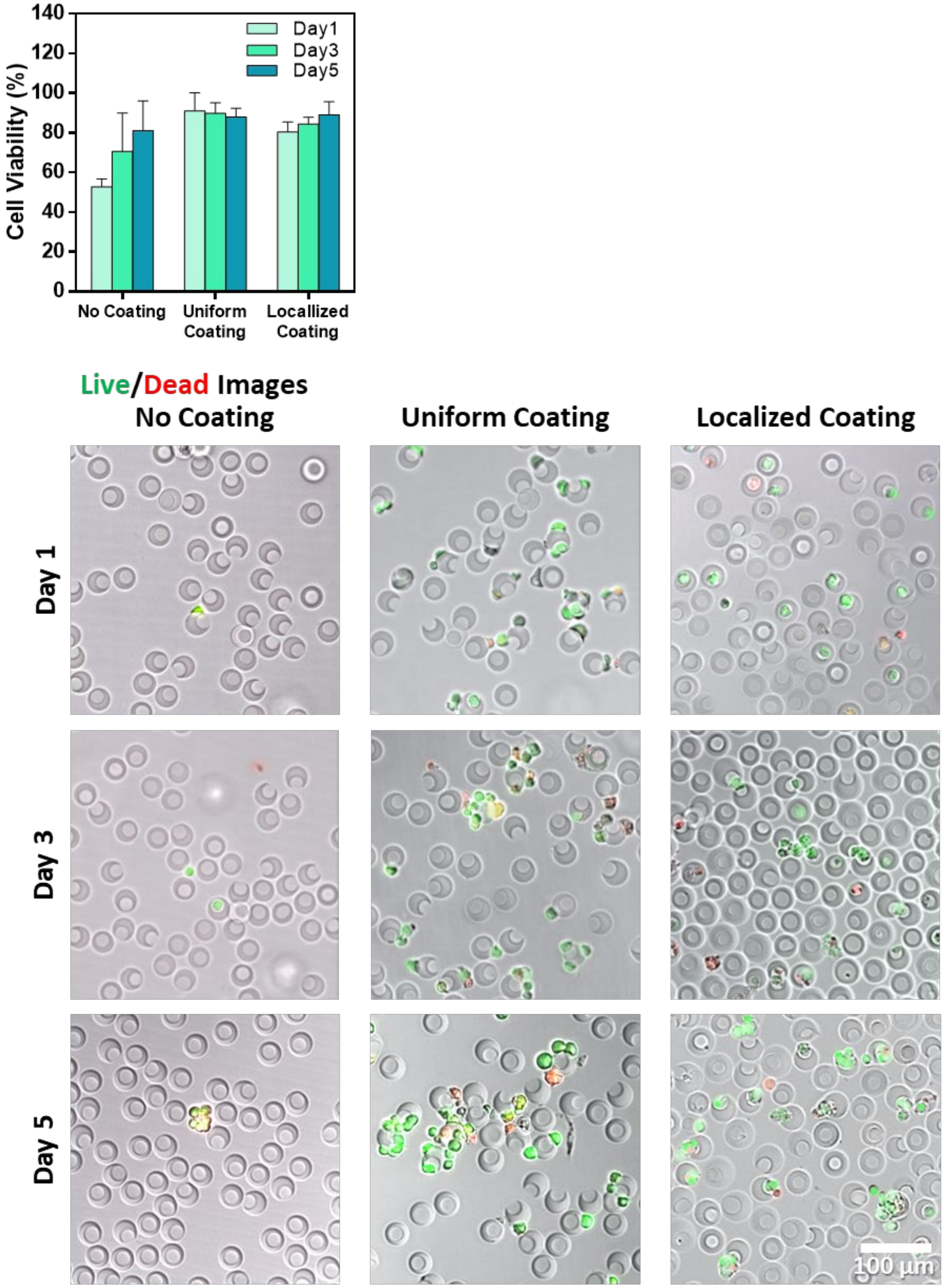


Figure S7. Viability of cells bound to nanovials with different cell binding moieties as reported in Figure 3. No Significant difference in viability was observed for the samples with uniform and localized coating of binding moieties.

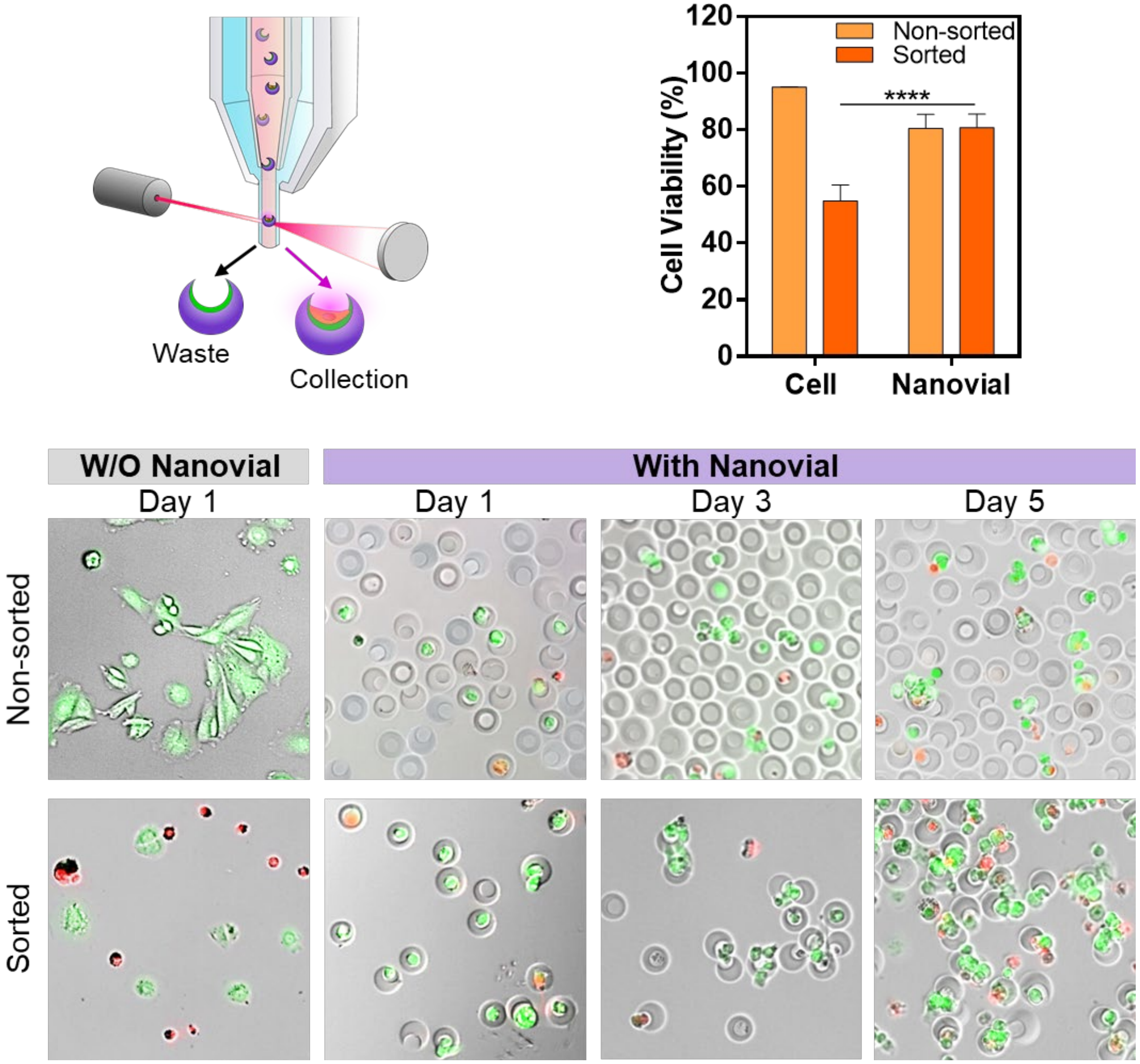


Figure S8. Cells encapsulated in nanovials showed significantly higher cell viability as compared to unbound cells after sorting. Nanovials presumably protect cells from hydrodynamic shear stress during the sorting process.

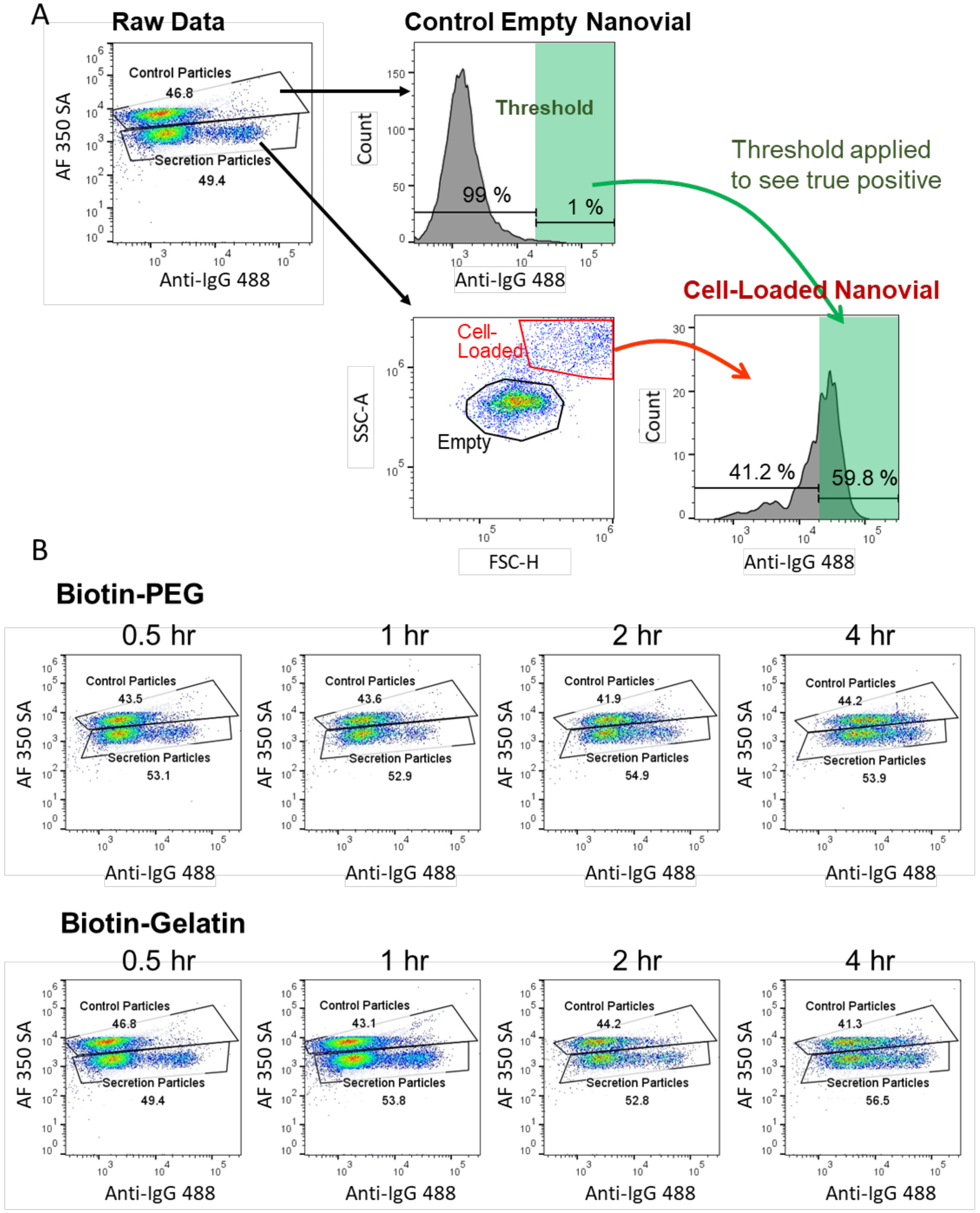


Figure S9. Flow cytometry gating strategy and scatter plots for secretion assay samples from Figure 6. (A) Cells that fall off during the assay can potentially skew the higher false positive signal. To account for this variability particles stained with AF350 were spiked into the population at the beginning of the secretion assay to quantify amount of cross-talk signal. This population was first gated from the remainder of the population to define as our empty nanovial intensity. The remaining population was gated and the cell containing sub-population was then gated based off increased forward and side scatter signal. (B) Raw scatter plots and gating for all of the samples.

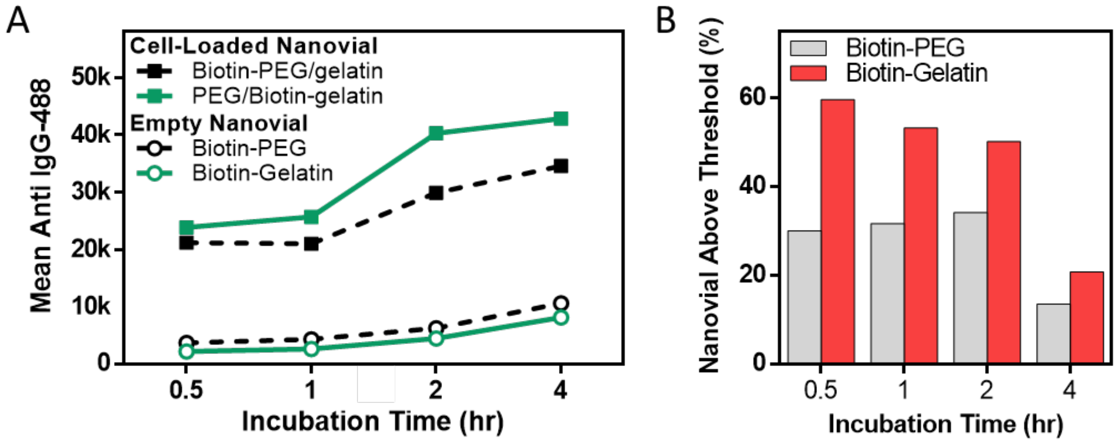


Figure S10. Additional characterization of the single cell secretion assay shown in figure 5. (A) Biotin-Gelatin nanovials showed lower IgG secretion signal on empty nanovials, while having higher signal on cell-laden nanovials when compared to the same assay performed on biotin-PEG nanovials. (B) When we defined a threshold of fluorescence intensity to have an only 1 % false positive rate, Biotin-Gelatin nanovials had a higher fraction of nanovials with positive signal above this threshold, indicating a significant reduction in cross-talk. For both nanovials, the fraction dropped as the nanovials were incubated longer than 2 hours.

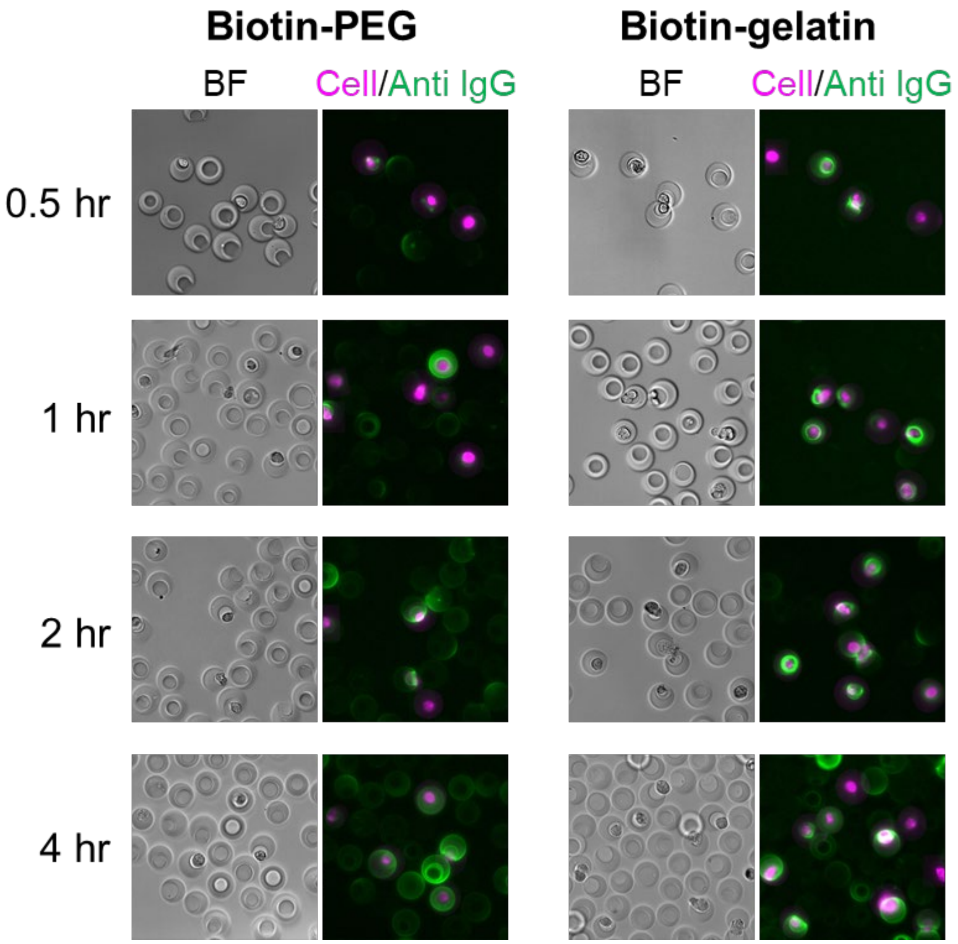


Figure S11. Brightfield and fluorescence microscopy images of nanovials and cells after an IgG secretion assay and prior to sorting. Biotin-Gelatin nanovials have signal more concentrated to the cavity and only start showing cross-talk at later time points. Signal on the biotin-PEG nanovials tends to bind more uniformly across the nanovial surface and shows signal on empty nanovials at earlier timepoints.
